# Directing Cell Delivery to Murine Atherosclerotic Aortic Lesions via Targeting Inflamed Circulatory Interface using Nanocarriers

**DOI:** 10.1101/2024.02.02.578719

**Authors:** Carlos Theodore Huerta, Leiming Zhang, Yulexi Y. Ortiz, Yan Li, Elnaz Zeynaloo, Emre Dikici, Teruna J. Siahaan, Sapna K. Deo, Sylvia Daunert, Zhao-Jun Liu, Omaida C. Velazquez

## Abstract

Stem cell therapy holds significant potential for many inflammatory diseases and regenerative medicine applications. However, delivery of therapeutic cells to specific disease sites after systemic administration without indiscriminate trafficking to other non-target tissues is a major limitation of current cell therapies. Here, we describe a novel nanocarrier-directed targeted cell delivery system that enables cell surface coating with dendrimer nanocarriers containing adhesion moieties to serve as a global positioning system “GPS” to guide circulating cells to targeted lesions and mediate the anchoring of cells at the inflammation site. By exploiting cell surface ligands/receptors selectively and/or molecular moieties that are highly expressed on activated endothelium in pathologic disease states, nanocarrier-coated cells containing the counterpart binding receptors/ligands can be enabled to specifically traffic to and dock at vasculature within target lesions. We demonstrate the efficacy of the I-domain fragment of LFA-1 (*id*LFA-1) complexed to modified nanocarriers to facilitate homing of mesenchymal stem cells (MSCs) to inflamed luminal endothelial cells on which ICAM-1 is highly expressed in a murine model of aortic atherosclerosis. Our method can overcome challenges imposed by the high velocity and dynamic circulatory flow of the aorta to successfully deliver MSCs to atherosclerotic regions and allow for docking of the potentially therapeutic and immunomodulating cells. This targeted cell-delivery platform can be tailored for selective systemic delivery of various types of therapeutic cells to different disease areas.

## 1. INTRODUCTION

The effectiveness of stem cell therapy is contingent upon effective cellular engraftment and homing to diseased tissues in order to reestablish function and homeostasis. Administration of stem cells is most commonly accomplished via local injection to the site of disease or intravascular delivery, including intraarterial injection that quickly delivers therapeutic cells to the tissue/organs fed by a given artery, and intravenous routes, by which therapeutic cells are infused into bloodstream (systemic delivery).^1,2^ However, these delivery routes are associated with substantial limitations. The most commonly employed administration route is direct injection, but cell-based therapy is significantly limited by cell viability and retention even when locally dispensed to sites of disease.^1,3^ Furthermore, cells delivered via local injection may not function appropriately given environmental factors at the tissue level including hypoxia, poor blood flow, hyperglycemia, and wide-spread local inflammation associated with disease.^4–7^ Limited space and even physical pressure at sites of inoculation may further change cellular characteristics or hamper cell viability.^5,8,9^ Unfortunately, many disease sites are not amenable to access by local inoculation given their intra-cavitary location (brain, chest, abdomen, aorta, etc.), which would necessitate more invasive injection procedures. Systemic administration via intravenous dissemination can partially overcome this limitation in delivery to sites that are traditionally hard to access; however, it can lead to indiscriminate trafficking of cells and often results in a low number of cells delivered to diseased tissue.^1,10^ Consequently, there is a critical, unmet need for novel methods to enhance the delivery of sufficient number of therapeutic cells to diseased tissue sites.

At the circulatory system interface, endothelial cells form a selectively permeable luminal barrier between blood and the neighboring tissue in homeostatic conditions. After physiological insult from tissue injury, malignancy, and/or inflammation, the release of numerous chemokines and cytokines such as TGF-β and SDF1-α activates neighboring endothelial cells (ECs).^11,12^ Resultant stimulation by these soluble factors can cause a variety of cell adhesion molecules (CAMs) such as integrins and selectins to be upregulated within the endothelium in the diseased tissue; thereby, making the endothelial lining “sticky” and increasing its ability to tether circulating cells responding to tissue repair, inflammation, and immunomodulation signals.^12^

Intercellular adhesion molecule (ICAM)-1 is a transmembrane cell surface molecule expressed by multiple cell types such as ECs. ICAM-1 plays essential roles in both innate and adaptive immune responses, as well as recruitment of circulating leukocytes for trans-endothelial migration and inflammation. While ICAM-1 is constitutively expressed at a low basal level on ECs, its levels are augmented by inflammatory cytokines (TNF-α, IL-1, and IFN-γ) in diseased states such as atherosclerosis. Atherosclerosis-associated cardiovascular diseases (CVDs) are the leading cause of morbidity and mortality globally and claim the lives of over 17 million people annually.^13–15^ Previously thought to be a bland lipid storage disease, atherosclerosis is now more appropriately characterized by its chronic inflammatory nature.^16^ Moreover, MSCs exhibit noteworthy anti-inflammatory properties^17,18^, providing new options for the treatment of inflammatory diseases. Hence, targeted delivery of exogenous therapeutic MSCs to atherosclerotic lesions can be a potential therapeutic option.

Crucial interactions between immune cells and orchestrated CAM levels on arterial ECs such as ICAM-1 are thought to be an early event in the natural history of atherosclerotic lesions, as well as in the migration of such cells to subendothelial segments and development into foam cells.^19,20^ Several ligands are able to bind to ICAM-1 of which the predominant counter-receptor mediating leukocyte interaction and recruitment is leukocyte function-associated antigen (LFA)-1. Specifically, the α-subunit of LFA-1 includes an amino-terminus, I-domain, which is responsible for LFA-1 binding to ICAM-1.^21–24^ ICAM-1 signaling transduction further contributes to the proinflammatory milieu and activation of ECs in blood vessel walls which propagate atherosclerotic plaque development.^19,25^ Preclinical evidence has shown significant upregulation of ICAM-1 at atherosclerotic-prone sites in murine models of cardiovascular disease that are homozygous apolipoprotein E-deficient (ApoE^-/-^), particularly within the aorta.^25,26^ Conversely, deficiency of ICAM-1 is protective against atherosclerosis and associated with substantially diminished atheromatous plaque burden *in vivo*.^27–29^ Consequently, the interaction between the I-domain of LFA-1 (*id*LFA-1) and ICAM-1 serves as an attractive therapeutic target for atherosclerotic disease of the circulatory system.

Herein we describe a novel application of nanocarriers with programmed molecular moieties based on the interaction of *id*LFA-1/ICAM-1 in order to control the localization and delivery of therapeutic stem cells to target disease regions. Specifically, fifth generation poly(amidoamine) (PAMAM) dendrimer nanoparticles were complexed with an adhesive moiety, *id*LFA-1, to form nanocarriers. In turn, these were utilized to install *id*LFA-1 on the surface of MSCs, which normally do not express LFA-1. Given our data and the previously mentioned expression pattern of ICAM-1 in luminal ECs at areas of atherosclerosis, *id*LFA-1 was selected as a targeting signal to enhance the docking and capture of the systemically delivered cells on the endothelial lumen at regions of vascular vessels expressing elevated levels of ICAM-1.

Dendrimer nanoparticles complexed with the adhesion molecule of interest were physically incorporated on the cell surface by complexation via ionic interactions between positively charged dendrimer nanoparticles and negatively charged cell membranes of MSCs. Furthermore, PAMAM dendrimers were optimized by acetylation to minimize intracellular internalization and toxicity and maximize cell surface coating as previously described.^5^ MSCs coated with *id*LFA-1 modified nanocarriers can be then recruited to target tissue sites expressing the cognate binding ligand (in this case ICAM-1 on endothelial lumen in atherosclerotic lesions). In our work we demonstrated the ability of both the target protein moiety and modified nanocarriers to bind ECs *in vitro* followed by the efficient, selective delivery of nanocarrier coated MSCs *in vivo* to atherosclerotic lesions in the aorta, thereby reducing indiscriminate migration to other organs in the ApoE^-/-^ mouse model of atherosclerosis. Hence, this nanocarrier technology provides a highly targeted biocompatible strategy to augment tissue homing of MSCs that can be applied to numerous other therapies ranging from cardiovascular and autoimmune diseases to regenerative medicine applications.

## 2. METHODS

### Reagents

Magnesium chloride, magnesium sulfate, sodium phosphate dibasic, potassium phosphate monobasic, sodium chloride, potassium chloride, ethylene diamine tetraacetic acid (EDTA), Tris−HCl, Tween 20, benzamidine hydrochloride, glycerol, ampicillin and sodium bicarbonate were purchased from Sigma-Aldrich (St. Louis, MO). VivoTag XL 680 NHS ester was purchased from Perkin Elmer. Protein-Free blocking buffer, Dithiothreitol (DTT), isopropyl-β-D-thiogalactopyranoside (IPTG), ProBlock GOLD protease inhibitor, and guanidine hydrochloride were obtained from G Biosciences (St. Louis, MO). Pierce 1-Step Ultra TMB-ELISA, Pierce BCA Protein Assay kit, *E. coli* BL21 cells, LB Broth and the OPTIMEM media were acquired from Thermo Fisher Scientific (Waltham, MA). Primary antibody Mouse IgG1 anti-human CD11a was purchased from Biolegend (San Diego, CA). Secondary antibody was horseradish peroxidase (HRP) conjugated anti-mouse IgG antibody and was procured from Southern Biotech (Birmingham AL). KPL TMB Stop Solution was purchased from Sera Care (Milford, MA). 20% acetylated, Generation 5 poly(amidoamine) (PAMAM) dendrimer was purchased from 21st Century Biochemicals (Marlborough, MA)

### Apparatus

Q500 Sonicator Ultrasonic processor was purchased from QSONICA, (Newtown, CT). Milli-Q water was produced by using a Pure Lab FLEX II Water purifier from ELGA Water Systems (High Wycomb, United Kingdom). KrosFlo KR21 tangential flow filtration (TFF) equipment was purchased from Repligen (Waltham, MA). 96-well, clear bottom, high-binding polystyrene microtiter plates, and Slide-A-Lyzer® dialysis cassettes was from Thermo Fisher Scientific (Waltham, MA). 4-20% Tris-Glycine SDS-Page gels was purchased from Bio-Rad (Hercules, CA). Novex IEF Gel pH, 3-10 were from Thermo Fisher Scientific (Waltham, MA). The wash steps were performed using Molecular Devices (Sunnyvale, CA) MultiWash+ Plate washer using five cycles of 250 μL/well of wash buffer, employing a 10 s shaking step at the end of each cycle. Absorbance measurements were performed either on a Spectromax 190 microtiter plater reader from Molecular Devices (San Jose, CA), or a Clariostar Optima UV/Vis Spectrophotometer from BMB Labtech (Ortenberg, Germany).

### Animal models

Male ApoE homozygous knockout mice (strain B6.129P2-*APOE^TM1UNC^*/J; ApoE^-/-^) were obtained from Jackson Laboratory (JAX stock #002052). The mice were maintained and bred under standard pathogen-free conditions. All animal experiments described were approved by the University of Miami Institutional Animal Care and Use Committee (IACUC) under Protocol 22-090. Atherosclerosis model were prepared by feeding ApoE^-/-^ mice a high-fat diet (HFD: 0.2 % total cholesterol, saturated fat > 60% total fat, and high sucrose) (TD.88137 Envigo) for 18 weeks after weaning. To perform endpoint examination experiments, euthanasia was performed via inhaled CO_2_ followed by cervical dislocation post-CO_2_ to ensure proper euthanasia.

### Fluorescence and bioluminescence IVIS Imaging

To track the location and biodistribution of infused *id*LFA-1 as well as *id*LFA-1-nanocarrier-coated murine MSCs, we conducted mouse whole-body *IVIS* (In Vivo Imaging System) scanning. To test infused *id*LFA-1, HFD-fed ApoE^-/-^ mice at 22 weeks old were separated into four groups (n=5). Vivo-Tag 680 XL (PerkinElmer #NEV11119) was used as a fluorescent dye to track *id*LFA1 and control murine serum albumin (MSA, Lee Biosolutions #101-50) distribution.

Group 1: ApoE^-/-^ mice were injected intravenously via tail vein cannulation (*i.v.*) with 50 µg/mouse of *id*LFA-1-Vivo-Tag 680 XL; Group 2: ApoE^-/-^ mice were injected *i.v.* with 140 µg/mouse of MSA-Vivo-Tag 680 XL: Group 3: ApoE^-/-^ mice were injected *i.v.* with an equivalent volume phosphate buffered saline (PBS); and Group 4: ApoE^-/-^ mice were injected *i.v.* with VivoTag 680XL only. Thirty minutes post-infusion of various *id*LFA-1, MSA or PBS, mice were anesthetized with 2.5% isoflurane according to our institutional animal protocol and underwent whole-body scan using the Perkin Elmer Xenogen IVIS Spectrum system. After whole-body scanning, the mice were euthanized, and the aortas were harvested immediately for imaging and scanned using the IVIS Spectrum system.

Similarly, to test infused *id*LFA-1-nanocarrier coated mouse MSCs, mouse MSCs pre-transduced with Luc2^+^/Lentivirus (Luc2^+^/MSCs) were coated or uncoated and injected into four groups of HFD-fed ApoE-/-mice: Group1 *id*LFA-1-nanocarrier coated MSCs; Group 2: MSA-nanocarrier-coated MSCs; Group 3: Dendrimer-coated MSCs; Group 4: MSCs alone.

Fifteen minutes post-infusion via tail vein injection of various MSCs, mice were injected intraperitoneally (*i.p.*) with D-luciferin (150 m/kg). After an additional 15 minutes, mice were anesthetized with 2.5% isoflurane according to our institutional animal protocol and underwent whole-body scan using the Perkin Elmer Xenogen IVIS Spectrum system. After whole-body scanning, the mice were euthanized, and the aortas were harvested immediately for imaging and scanned using the IVIS Spectrum system.

Photon counts of either fluorescence and bioluminescence signals were acquired and analyzed using Living Image version 4.7 software. Regions of interest (ROIs) were drawn sections in the defined area and quantified using the physical, calibrated unit “Radiant Efficiency [p/sec/cm^²^/sr] / [µW/cm^²^]”. The Living Image software normalizes automatically for sensitivity differences resulting from different exposure times without any user input required when ROI values are expressed in a calibrated, physical unit.

### Preparation for recombinant *id*LFA-1 and nanocarriers

The expression of the *id*LFA-1 has been performed, with minor modifications, as described. ^30^ Briefly, a plasmid containing the I-Domain DNA sequence, pET-11a/LFA-1 was transformed into competent *E. coli* BL21 cells. The cells were grown in 5.0 mL LB Broth containing 100 µg/mL ampicillin. Next day, 300 mL of LB Broth containing 100 µg/mL ampicillin was inoculated with 5.0 mL of the refreshed overnight cultures. To refresh, the overnight cultures were centrifuged at 5,000 xg for 10 minutes, the spent LB broth was discarded, and the cell pellet was resuspended in the same volume of fresh LB broth. The flasks were incubated at 37 °C, with shaking at 250 rpm, until the optical density of the culture at 600 nm (OD600) was around 0.8. Then, 300 µL of 1.0 M IPTG and 3.0 mL of 1.0 M sterile magnesium chloride was added to have final concentrations of 1.0 mM and 10.0 mM, respectively. The flask was incubated at room temperature for 3.0 h, with shaking at 250 rpm. The culture was then centrifuged at 8,000 xg for 15 minutes at 4 °C to harvest the bacterial cells expressing *id*LFA-1 in the form of inclusion bodies. The LB broth was discarded, and the cell pellet was resuspended in 30 mL of homogenization buffer (50 mM TRIS buffer containing 1.0 mM DTT, 10 mM magnesium chloride, 2.0 mM EDTA, and 5.0 mM benzamidine hydrochloride at pH 8.00) and sonicated using 1 s pulses for 20 minutes at a 20% amplitude. After the sonication is complete, the cell suspension was centrifuged at 18,000 xg at 4 °C for 15 minutes. Then, the pellet was washed by resuspending in 30 mL of homogenization buffer and centrifuging at 18,000 xg at 4 °C for 15 minutes. This washing step was performed once more, followed by a final wash using deionized water. The pellet, which is consisting of denatured *id*LFA-1 in the form of inclusion bodies, was solubilized in about 30 mL of denaturation buffer (50 mM TRIS buffer containing 2.0 mM DTT, and 6 M guanidine hydrochloride at pH 8.00) by rotating at room temperature for 1.0 h. Then, the solution was centrifuged at 18,000 xg at 4 °C for 15 minutes to remove any insoluble matter. The solution was diluted in 200 mL cold DI water at 4 °C followed by rapidly diluting in 4.0 L of cold renaturation buffer (50 mM TRIS buffer containing 1.0 mM DTT, and 5 mM magnesium sulfate and 7% glycerol at pH 8.00) to refold the *id*LFA-1 protein into its functional form. This solution contains dilute, purified, *id*LFA-1 which was then concentrated using 3,500 MWCO tangential flow filtration column. After concentrating the protein sample 40-fold, the sample was dialyzed using 10,000 MWCO dialysis cassettes into phosphate buffered saline at pH 7.4. Any formed precipitate during dialysis was removed by centrifuging at 18,000 xg at 4 °C for 15 minutes. The purity of the protein was confirmed using sodium dodecyl sulfate polyacrylamide gel electrophoresis (SDS-PAGE), followed by the measurement of the protein concentration using BCA assay. The isoelectric point of the purified protein was determined using Novex IEF Gel pH, 3-10 according to the manufacturer’s protocol.

### CD Spectroscopy Analysis

Samples of *id*LFA-1 were dialyzed into CD buffer containing 10 mM potassium phosphate and 100 mM ammonium sulfate at pH 7.40 at a final concentration of 0.1 mg/mL and the far-UV CD spectra were recorded using a Jasco J-815 Spectropolarimeter (Jasco, Easton, MD, USA) (Scan mode: continuous; scan speed 50 nm/min; data pitch 0.5 nm; bandwidth: 1 nm; data integration time (D.I.T.) 2 s; accumulations: 3).^31^ Analysis of the CD data was performed using Dichroweb (http://dichroweb.cryst.bbk.ac.uk) and secondary structure assignments were generated using the CDSSTR analysis package.

### Labeling of *id*LFA-1, MSA and BSA with fluorophores Cy5, and VivoTag XL 680 (VT-XL 680)

The proteins used in this study were labeled with the fluorophores Cy5 and VT-XL680 according to the manufacturer’s protocols. Briefly, an aliquot of 1.0 mg/mL of the protein, dissolved in 50 mM carbonate/bicarbonate buffer at pH 8.00, was reacted with the 4x mole excess of the fluorophore, dissolved at a concentration of 1.0 mg/mL in anhydrous DMSO, at room temperature for 2 h. The excess, unreacted fluorophore, was then removed by dialysis against PBS at pH 7.40, and the protein concentration and the fluorophore labeling efficiency was measured according to the manufacturer’s protocol using a UV-Vis spectrophotometer.

### Preparation of Tagged proteins-Ac-G5 nanocarriers and cell surface coating with nanocarriers

The nanocarriers were prepared by complexing 2x mole excess of commercially purchased 20% acetylated G5-PAMAM dendrimers with either labeled or unlabeled proteins. Briefly, in order to coat 1.0 x 10^6^ cells, 1.4 nanomoles of protein of interest and 2.8 nanomoles of 20% acetylated G5-PAMAM dendrimer was mixed at room temperature for 15 minutes. Both the dendrimer and the protein were dissolved in OptiMem medium separately and the dendrimer solution was added, dropwise, to the protein solution, while mixing. The mixture was then incubated at room temperature for 15 minutes to allow for the complexation. Any formed precipitate during incubation was removed by centrifuging at 18,000 xg at 4 °C for 15 minutes.

### Preparation of MSCs coated with Nanocarriers

To coat the MSCs, an aliquot of 1.0 mL of nanocarriers, prepared as described above, in OptiMem medium were mixed with 1.0 x 10^6^ MSCs and were incubated for 20 min at room temperature with gentle mixing every 5 minutes. Afterwards, nanocarrier coated MSCs were centrifuge at 270 xg for 5 minutes and were gently resuspended with a pipette in 5.0 mL sterile PBS.

### Recombinant lentiviruses and cell transduction

DsRed/Lentiviral vector plasmid, LacZ/Lentiviral vector plasmid and Luc2/Lentiviral vector plasmid were described previously.^5^ Human ICAM-1/Lentiviral vector was purchased from GenTarget Inc, (LVP595, San Diego, CA). Production of pseudo-typed lentivirus was achieved by co-transfecting 293 T cells with three plasmids as described.<u>^10^</u> The lentiviruses collected 48 hours post-transfection displayed titers of around 10^7^ transducing units/ml as determined by PCR. Mouse bone marrow-derived MSCs^32^ and human microvascular endothelial cells (HMVEC)^31^ were prepared and cultured as previously described. To infect cells by lentivirus, cells were exposed for six hours to virus with an MOI (multiplicity of infection) of 5 viral particles/cell in the presence of 4 μg/mL polybrene (Sigma-Aldrich). Cells were then washed, cultured with regular complete medium for two additional days, and analyzed by fluorescence microscopy (for DsRed^+^/MSCs) or immunoblotting (for ICAM-1/Lentivirus-transduced HMVEC^ICAM-1*hi*^ and LacZ/Lentivirus transduced HMVEC^ICAM-1*lo*^ or bioluminescence reader (SpectraMax L, Molecular Devices) (for Luc2^+^/MSCs), respectively, using standard protocols.^5^ Cells were pooled for subsequent analysis as indicated in individual experiments.

### *In vitro* binding assays

1 x 10^5^ cells/well of HMVEC^ICAM-1*hi*^ were cultured in the 24-well glass plates (P^11^24G-1.5-10-F, MatTek) pre-coated with 1% gelatin and cells reached 100% confluence one-day later. To test the binding of fluorescent dye-labelled *id*LFA-1 or *id*LFA-1-nanocarrier on HMVEC^ICAM-1*hi*^, 100 μM *id*LFA1-Cy5 *vs* BSA-Cy5, or *id*LFA1-Cy5-conjugated Ac-G5 nanocarrier *vs* BSA-Cy5-conjugated Ac-G5 nanocarrier were added to the well in which a HMVEC monolayer was formed. After incubation for 30 minutes at 37 °C, wells were washed with PBS twice, and Cy5 signals remained in wells were visualized by fluorescence microscopy and quantified by fluorescence scanner (GE Typhoon Trio, Piscataway, NJ). Similarly, to test binding *id*LFA-1-nanocarrier-coated cells on HMVEC^ICAM-1*hi*^, 1 x 10^5^ mouse bone-marrow-derived MSCs pre-transduced with DsRed/Lentivirus (DsRed^+^/MSCs), which were pre-coated with *id*LFA-1-Ac-G5 and BSA-Ac-G5 nanocarriers, and suspended in 1 mL basal MesenCult^TM^ (StemCell Technologies, Cambridge, MA) medium and added to the well in which a HMVEC monolayer was formed and incubated for 1 hour at 37°C. Unbounded MSCs were washed out twice with PBS. Red fluorescence signals derived from adherent DsRed^+^/MSCs were measured and quantified by fluorescence scanner. All *in vitro* binding assays were duplicates in 24-well plates and experiments were repeated three times.

### Immunofluorescence assays

Immunofluorescence studies were performed in tissue sections to evaluate expression of ICAM-1 on endothelial cells in aorta, homing of nanocarrier-coated MSCs to atherosclerotic lesions, and biodistribution of nanocarrier-coated MSCs in lung and liver. Tissue section slides were deparaffinized per standard protocol and antigen retrieval was performed in EDTA buffer (pH 9.0) at 100 °C for 10 minutes. Slides were then washed in distilled water and permeabilized with 0.25% Triton-X100 TBS for 15 minutes and rinsed twice with PBS. Slides were subsequently incubated with Protein Block (x0909, Agilent Dako) for 1 hour. To detect various proteins, the following antibodies were utilized: Alexa Fluor® 488 anti-CD31 antibody (EPR17259, ab305267, Abcam), Alexa Fluor® 647 anti-Firefly Luciferase antibody (EPR17790, ab233049, Abcam), Alexa Fluor® 488 E-Selectin/CD62E antibody (103), NBP289430AF488, Novus Biologicals. Slides were incubated overnight at 4 °C with antibodies in protein block for 1 hour at room temperature. The slides were then washed with 0.1% Tris-buffered saline and Tween-20 (TBST) prior to 4’,6-diamidino-2-phenylindole dihydrochloride (DAPI) staining (D9542 Sigma) for nuclei visualization. Slides were imaged utilizing an Olympus IX71 Inverted fluorescence microscope.

### Statistical Analyses

Analysis of statistical differences were performed utilizing ANOVA for multiple samples and 2-tailed Student’s *t*-test for pairwise comparison. Data are expressed as mean ± standard deviation (SD), and values are considered significant based on a threshold of *p*<0.05.

## 3. RESULTS

### ICAM-1 expression is elevated on endothelium in aorta of ApoE^-/-^ mice with atherosclerosis

Atherosclerosis is a chronic vascular inflammatory disease. ICAM-1 is well known to play a role in the recruitment of immune cells. to sites of inflammation, including atherosclerotic lesions.

To establish the presence of ICAM-1 in atherosclerotic lesions in our experimental animal model, we conducted immunofluorescence analysis (IFA) to examine the expression of ICAM-1 on luminal endothelium in the aorta of 22-week old ApoE-/-mice fed with HFD to recapitulate atherosclerosis. IFA confirmed elevated expression of ICAM-1 colocalization with the luminal endothelial cell marker CD31 in aortas of ApoE^-/-^ mice fed with HFD+. Elevated levels of ICAM-1 were observed on luminal endothelium, small vessels and capillaries in tunica intima and adventitia (Figure 1A). In contrast, expression of ICAM-1 was rarely detectable in the aorta of control ApoE^-/-^ mice fed standard diets (HFD-). Interestingly, levels of ICAM-1 were robustly higher in endothelium in plaques (Figure 1B), indicating a correlation of vascular inflammation status and ICAM-1 expression. These results confirmed that ICAM-1 expression is elevated on inflamed endothelium in aortas, especially in plaques, of ApoE^-/-^ mice with atherosclerosis.

**Figure 1.**
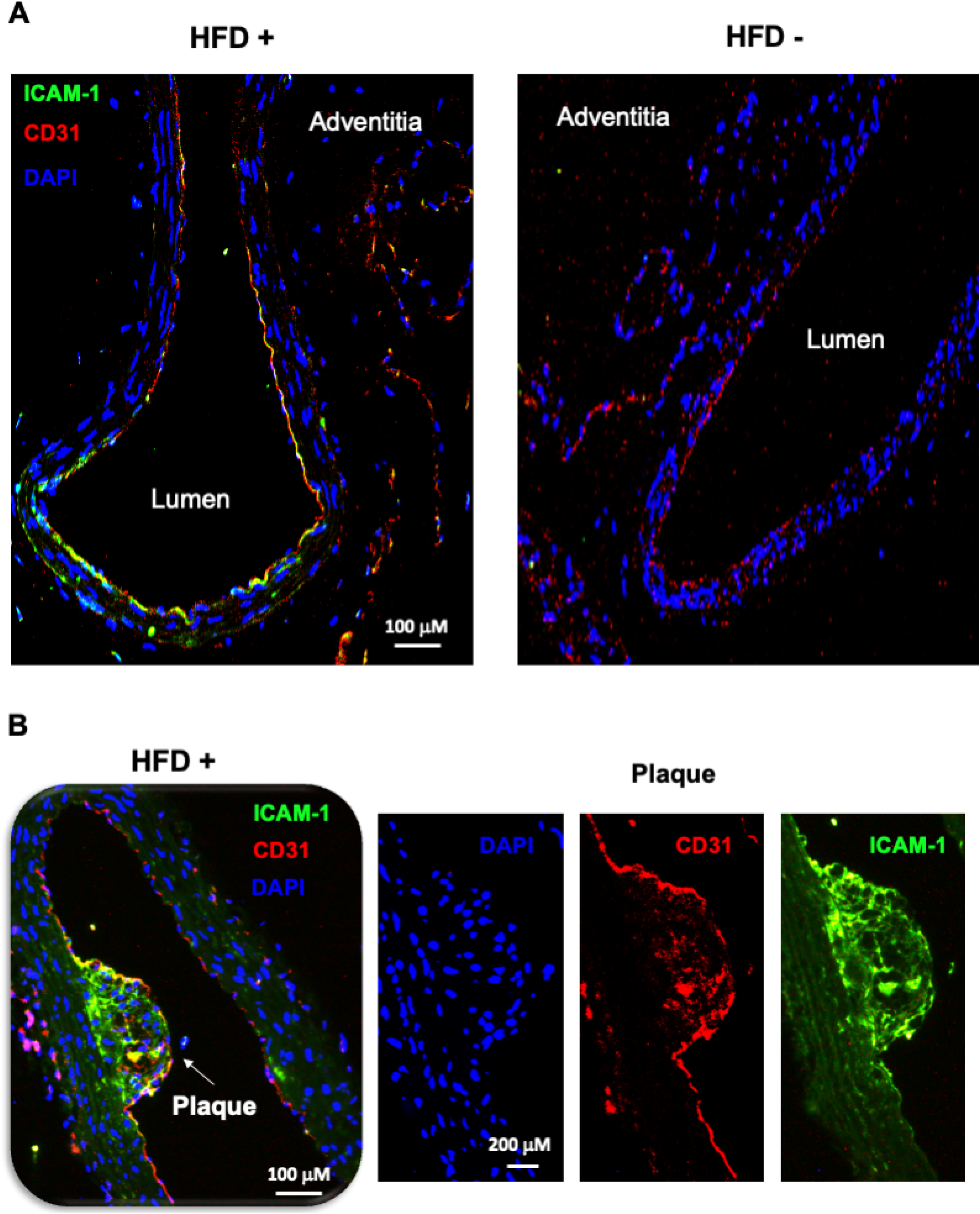
Increased expression of ICAM-1 in luminal endothelium of the aorta in ApoE^-/-^ mice. Representative images shown **c**o-expression (yellow) of ICAM-1 (green) and CD31 (red) in luminal endothelium at areas without plaques **A** and at plaques **B** within aortas from ApoE^-/-^ mice fed high-fat diet (HFD +) compared to standard diet (HFD -). Images of individual staining with CD31 (red), ICAM-1 (green) and DAPI (nucleus) at plaque are also shown in **B**.

### Purification of idLFA-1 and verification of idLFA-1/ICAM-1 association

We sought to take advantage of ICAM-1/LFA-1 adhesion molecule pairs to mediate cell-cell interaction in the circulatory interface to achieve targeted cell delivery (directly homing the infused circulating *id*LFA-1-coated MSCs to the ICAM-1-expressing inflamed ECs on the lumen of the aorta). Since ICAM-1 expression is elevated on endothelium in aorta of ApoE^-/-^ mice with atherosclerosis, we utilized the I-domain of LFA-1 (*id*LFA-1) as a payload to create nanocarriers given that I-domain (*id*) is responsible for mediating ICAM-1/LFA-1 interactions. In addition, using *id,* a smaller fragment of LFA-1, can increase the amount of payload in nanocarriers, yet decrease cost and potential side-effects caused by large/whole LFA-1 protein. Nanocarrier systems typically comprise nanoparticles (dendrimers, PLGA, liposomes) and payloads, such as the target agent or protein moiety, to form vehicles to guide the cells to the proper location. The *id* of LFA-1 was overexpressed and purified from *E. coli* cells, in the form of inclusion bodies.

The inclusion bodies have native-like secondary structure of the expressed protein, and they are resistant to proteolytic degradation. Since they are formed due to specific molecular interactions among a single type of protein, they mostly consist of the target recombinant protein in relatively high purity. ^33^ Therefore, our purification strategy to obtain pure *id*LFA-1 involved isolating the formed inclusion bodies of *id*LFA-1, extensive washing of the protein pellet, followed by a denaturation and renaturation cycle as described by Manikwar *et al*. ^24^ The purified protein was then characterized for its purity. In SDS-PAGE gel, the purified protein runs as a single band with an apparent molecular weight of 19.7 KDa (Figure 2A). The protein was also characterized with respect to its tertiary structure using circular dichroism (CD) spectroscopy. The purified *id*LFA-1 protein was consistent with the expected *id*LFA-1 characteristics (Figure 2B). To test binding activity of *id*LFA-1, we carried out a cell binding assay using *id*LFA-1 conjugated to the fluorescent probe Cy5 for visualization and quantification by fluorescence microscopy and scanner, using a previously described method.^5^

**Figure 2.**
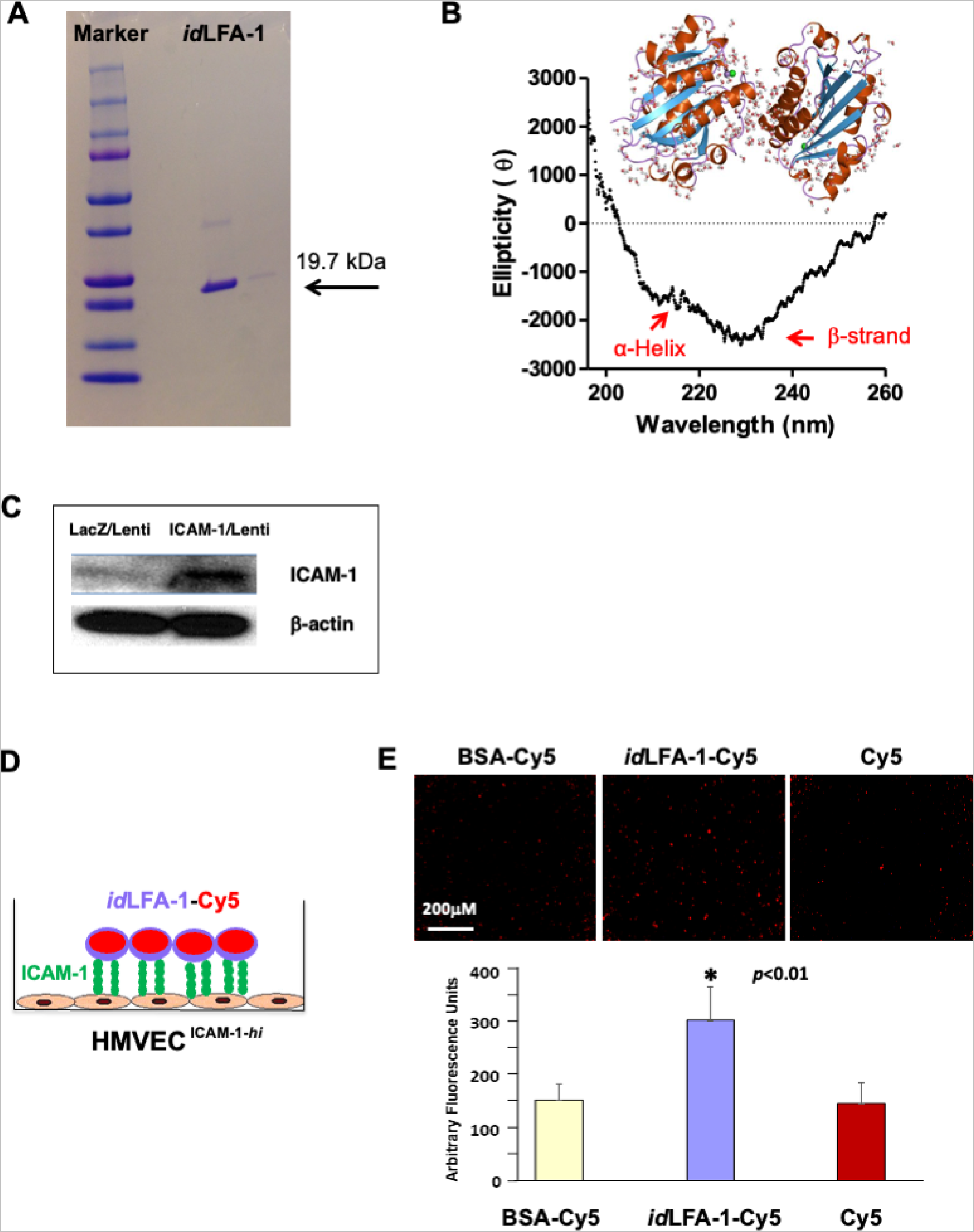
Verification of recombinant *id*LFA-1 purity and binding capability with ICAM-1 *in vitro*. **A**. Recombinant purification of *id*LFA-1 with the appropriate molecular weight (19.7 kDa) was verified. **B**. Circular Dichroism spectra were collected to confirm the appropriate tertiary structure (α-Helix and β-strand) of purified *id*LFA-1. **C**. Western blot analysis demonstrating overexpression of ICAM-1 protein in HMVEC transduced with ICAM-1/Lentivirus and control LacZ/Lentivirus. **D**. Schematic illustration of the *id*LFA-1-Cy5-(Ac-G5) dendrimer conjugate moiety binding to endothelial cells stimulated to express ICAM-1 (EC^ICAM-1-*hi*^). **E**. Microscopy imaging demonstrating association of *id*LFA-1-Cy5 with EC^ICAM-1-*hi*^ compared to BSA-Cy5-(Ac-G5) and Cy5-(Ac-G5) alone as well as quantitative data of Cy5 signal. Data are presented as mean ± SD of three independent assays in which samples were duplicated. Asterisk denotes statistical significance (*p*<0.01).

Overexpression of ICAM-1 was achieved by transduction of HMVEC with ICAM-1/Lentivirus vs LacZ/Lentivirus (control). Levels of ICAM-1 in HMVEC^ICAM-1*hi*^ *vs* HMVEC^ICAM-1*lo*^ was confirmed by immunoblotting (Figure 2C). Figure 2D illustrates how this in vitro binding assay is performed. HMVEC^ICAM-1*hi*^ were cultured in 24-well glass plates to form a cell monolayer. A concentration of 100 μM *id*LFA1-Cy5 (test) *vs* BSA-Cy5 (control) *vs* Cy5 (blank) were added to wells in which monolayers of HMVEC^ICAM-1*hi*^ had been established. After incubation for 30 minutes at 37 °C, wells were washed with PBS twice, and Cy5 signals remaining in wells were visualized by microscopy and quantified by fluorescence scanner. We observed significantly higher Cy5 signal remained in wells added with *id*LFA1-Cy5 compared to wells exposed to BSA-Cy5 and Cy5 alone (Figure 2E). These results demonstrated that purified *id*LFA1fragments carry the anticipated biological activity and can bind with ICAM-1 on the cell surface.

### Binding of idLFA-1-nanocarriers and idLFA-1-nanocarrier-coated MSCs with ICAM-1 in vitro

We next sought to validate the binding capability between *id*LFA-1-Cy5-(Ac-G5) dendrimer nanocarriers and endothelial cells expressing ICAM-1. We specifically investigated the interaction between *id*LFA-1-Cy5-(Ac-G5) and HMVEC^ICAM-1*hi*^. For this, the *id*LFA-1-Cy5 was complexed to Ac-G5 dendrimer to form nanocarriers (*id*LFA-1-Cy5-(Ac-G5)). BSA nanocarriers (BSA-Cy5-(Ac-G5)) were constructed as control. The *id*LFA-1-Cy5-(Ac-G5)) and BSA-Cy5-(Ac-G5)) were added to wells in which monolayers of HMVEC^ICAM-1*hi*^ were present (Figure 3A, *left*). After incubation for 30 minutes at 37 °C, wells were washed with PBS twice, and remaining Cy5 signals in wells were visualized by fluorescence microscopy and quantified by employing a fluorescence scanner. Through these experiments and by employing fluorescent microscopy, we found that *id*LFA-1-Cy5-(Ac-G5) exhibited superior adhesion to the HMVEC^ICAM-1*hi*^ monolayers compared to BSA-Cy5-(Ac-G5) (Figure 3A, *right*). This further demonstrated that *id*LFA-1-nanocarriers are able to bind to ICAM-1 on the surface of ECs.

**Figure 3.**
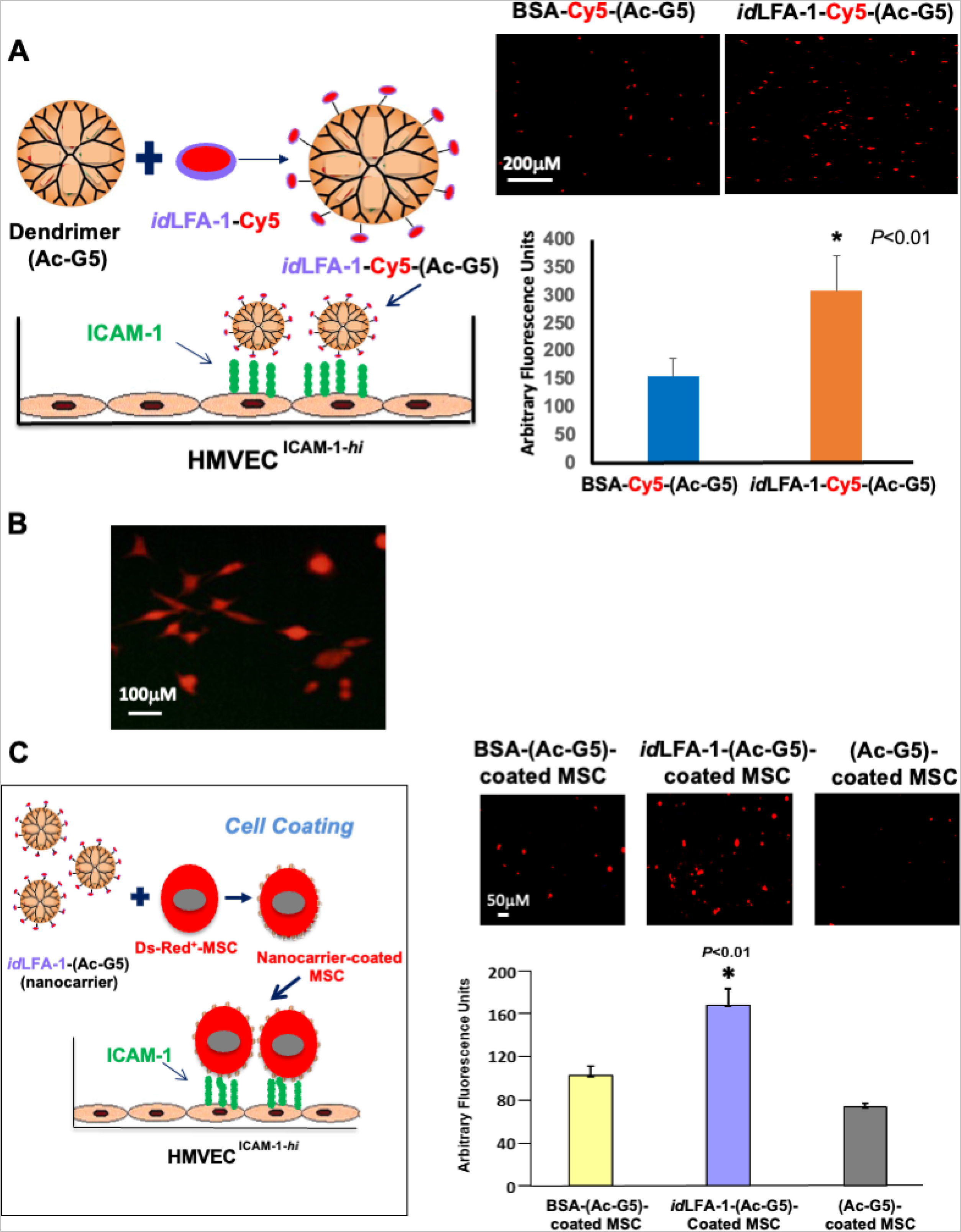
*id*LFA-1-Cy5-(Ac-G5) dendrimer nanocarriers and *id*LFA-1-Cy5-(Ac-G5) nanocarrier-coated MSCs preferentially bind to ICAM-1 expressing endothelial cells *in vitro*. **A**. *Left*: Schematic illustration of the *id*LFA-1-Cy5-(Ac-G5) dendrimer conjugate moiety binding to HMVEC transduced with ICAM-1/Lentivirus to overexpress ICAM-1 (HMVEC^ICAM-1-*hi*^). *Right*: Fluorescent microscopy imaging demonstrating increased association of *id*LFA-1-Cy5-(Ac-G5) with HMVEC^ICAM-1-*hi*^ compared to BSA-Cy5-(Ac-G5). **B**. Fluorescent microscopy demonstrating Ds-Red fluorescent expression in MSCs after transduction with Ds-Red/Lentivirus. **C**. *Left*: Schematic illustration of binding of *id*LFA-1-Cy5-(Ac-G5)-coated MSCs with HMVEC^ICAM-1-*hi*^. *Right*: Fluorescent microscopy imaging demonstrating association of *id*LFA-1-(Ac-G5)-coated MSCs binding with HMVEC^ICAM-1-*hi*^ compared BSA-nanocarrier and nanocarrier alone coated MSCs. Quantitative data of Cy5 signal. Data are presented as mean ± SD of three independent assay in which samples were duplicated. Asterisks denotes statistical significance (* *p*<0.01).

We next tested whether *id*LFA-1-nanocarrier-coated MSCs were able to bind to ICAM-1 on the surface of ECs. Mouse bone-marrow-derived MSCs, pre-transduced with DsRed/lentivirus (DsRed^+^/MSCs) (Figure 3B), were coated with *id*LFA-1-(Ac-G5) nanocarriers and BSA-Cy5-(Ac-G5)- and Cy5-(Ac-G5) nanocarriers-coated MSCs (controls). Similarly, three nanocarrier-coated MSC groups (*id*LFA-1-Cy5-(Ac-G5)-, BSA-Cy5-(Ac-G5)- and Cy5-(Ac-G5)-coated MSCs) were added to wells in which monolayers of HMVEC^ICAM-1*hi*^ were seeded as illustrated in Figure 3C, *left*. After incubation for 30 minutes at 37 °C, wells were washed with PBS twice, and DsRed^+^/MSCs associated with HMVEC^ICAM-1*hi*^ in wells were visualized by fluorescence microscopy and quantified by using a fluorescence scanner. We determined that there were significantly higher numbers of *id*LFA-1-Cy5-(Ac-G5)-coated MSCs associated with HMVEC^ICAM-1*hi*^ compared to BSA-Cy5-(Ac-G5)- and Cy5-(Ac-G5) nanocarrier-coated MSCs (Figure 3C, *right*). Taken together, our data demonstrated that *id*LFA-1 is able to mediate cell— cell interaction in an *in vitro* model.

### idLFA-1 binds to atherosclerotic lesions in aorta of ApoE^-/-^ mouse

We then tested the ability of *id*LFA-1 to bind to inflamed atherosclerotic lesions in the aorta of an ApoE^-/-^ mouse fed with a HFD. The *id*LFA-1 was conjugated to a fluorochrome (Vivo-Tag XL680, PerkinElmer #NEV11119) for *in vivo* IVIS imaging. A volume of 100 μl of Vivo-tag-*id*LFA-1, Vivo-tag-MSA (murine serum albumin, MSA, Lee Biosolutions #101-50) and Vivo-tag alone were intravascularly injected to 22 weeks old ApoE^-/-^ mouse fed HFD via tail vein (*i.v.*) (5 mice/group). Development of atherosclerosis in the aorta of ApoE^-/-^ mice was pre-demonstrated by *Oil Red O* staining (Figure 4A). Mice were subjected to whole-body IVIS scan thirty minutes following *i.v.* injection. Compared to control mice injected with Vivo-tag-MSA (1.19 ± 0.84 x 10^9^ [p/s/cm²/sr] / [µW/cm²]) and Vivo-tag alone (4.42 ± 4.80 x 10^8^ [p/s/cm²/sr] / [µW/cm²]), a significantly higher amount of Vivo-tag-*id*LFA-1 was present in the region of the aorta (2.82 ± 0.28 x 10^9^ [p/s/cm²/sr] / [µW/cm²] (Figure 4B). Vivo-tag signals in bladders of all three groups of mice are comparable, indicating that equal amounts of Vivo-tag-*id*LFA-1, Vivo-tag-MSA and Vivo-tag were injected into mice. The signals in the region of the aorta were normalized by that in the bladder. After whole-body scan, aortas were immediately resected and subjected to IVIS scan. Consistent with results of whole-body scan, a significantly higher amount of Vivo-tag-*id*LFA-1 signal was detected in the aorta of mice injected with Vivo-tag-*id*LFA-1 (1.06 ± 0.50 x 10^9^ [p/s/cm²/sr] / [µW/cm²]) compared to control mice injected with Vivo-tag-MSA (4.91 ± 1.77 x 10^8^ [p/s/cm²/sr] / [µW/cm²]) and Vivo-tag alone (5.47 ± 1.74 x 10^8^ [p/s/cm²/sr] / [µW/cm²]) (Figure 4C). Because Vivo-tag-*id*LFA-1 signals were richer in arch regions of aortas where mice have more severe atherosclerotic lesions, we also compared the intensity of the Vivo-tag-*id*LFA-1 signals in arch regions. Consistent with results of whole aorta, a robustly increased Vivo-tag-*id*LFA-1 signal intensity was presented in the arch of aorta in mice injected with Vivo-tag-*id*LFA-1 (3.99 ± 2.50 x 10^8^ [p/s/cm²/sr] / [µW/cm²]) compared to control mice injected with Vivo-tag-MSA (6.57 ± 6.02 x 10^7^ [p/s/cm²/sr] / [µW/cm²]) and Vivo-tag alone (4.66 ± 6.61 x 10^7^ [p/s/cm²/sr] / [µW/cm²]). Taken together, these results demonstrated that infused circulating *id*LFA-1 is able to home and bind to atherosclerotic lesions in aorta of ApoE^-/-^ mouse, indicating that *id*LFA-1 can be used as targeting moiety to create nanocarriers to mediate targeted cell delivery to atherosclerotic lesions *in vivo*.

**Figure 4.**
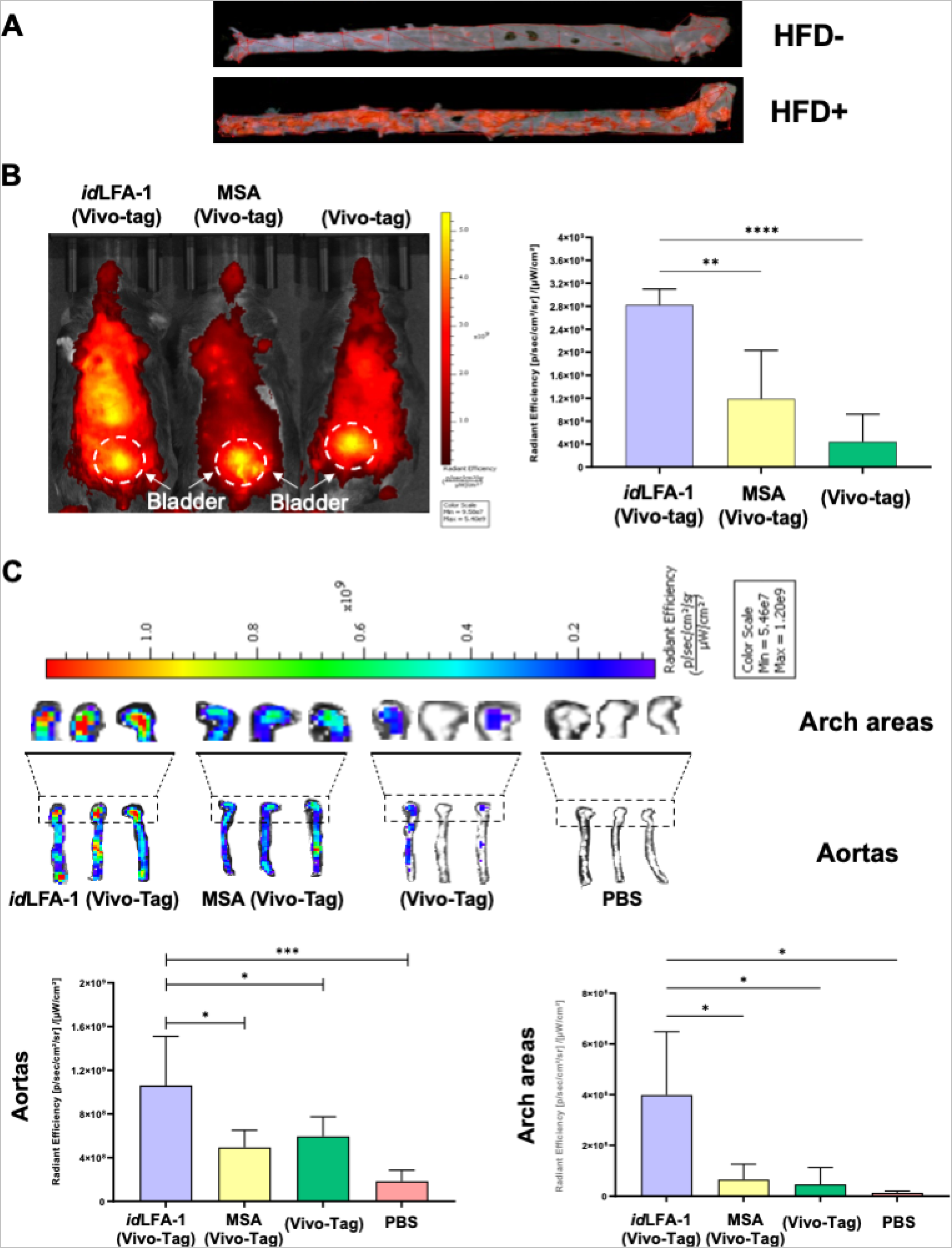
Targeted delivery of *id*LFA-1 to the aorta and aortic arch after systemic infusion *in vivo.* **A**. Whole aorta visualization of atherosclerotic plaques by Oil Red O staining in ApoE^-/-^ mice fed with or without HFD. **B**. *Left*: *In vivo IVIS* imaging shows an increased signal intensity of *id*LFA-1 conjugated to Vivo-tag dye throughout the body and most dense in the midline compared to murine serum albumin (MSA)-Vivo-tag and Vivo-tag controls 30 minutes after systemic injection in ApoE^-/-^ mice fed HFD. *Right*: Quantitative data of Vivo-tag fluorescent dye intensity in each group of mice (n = 5 / group). Asterisks denote statistical significance (** *p* = 0.022; **** *p* <0.0001). **C**. *Top*: Three representative *IVIS* imaging shows an increased uptake of *id*LFA-1 conjugated to Vivo-tag in the aortic arch areas and aortas harvested from ApoE^-/-^ mice fed a HFD compared to MSA-Vivo-tag, Vivo-tag alone, and PBS controls. *Bottom*: Quantification data (ANOVA analysis) of fluorescent dye signal intensity demonstrates higher signal in the aorta and arch areas of mice receiving *id*LFA-1 conjugated to Vivo-tag. Asterisks denote statistical significance (* *p*<0.05; *** *p*<0.001, n=5/group).

### idLFA-1-nanocarrier-coated MSCs home to atherosclerotic aorta lesions in ApoE^-/-^ mouse model

Next, we employed *id*LFA-1 as a targeting moiety to create nanocarriers for targeted delivery of MSCs to atherosclerotic lesions *in vivo*. The *id*LFA-1-nanocarriers (*id*LFA-1-(Ac-G5)) and control nanocarriers (MSA-(Ac-G5)) were created and applied to coat mouse MSCs, which were pre-transduced with Luc2/Lentivirus as described above. A total of 1 x 10^6^ Luc2^+^/MSCs coated with *id*LFA-1-(Ac-G5) *vs* MSA-(Ac-G5) were suspended in 100 μl of PBS and intravascularly injected to 22-week old ApoE^-/-^ mouse fed with HFD via tail vein (*i.v.*) (5 mice/group). Mice injected with 100 μl of PBS were used as negative control. Fifteen minutes post cell infusion, mice were injected (*i.p.*) with D-luciferin. After an additional 15 min time elapsed, mice were sacrificed, and aortas (and major organs, include lung and liver, see biodistribution below), were harvested and subjected to IFA to detect homing of infused Luc2^+^/MSCs to atherosclerotic aorta lesions in ApoE^-/-^ mice. We observed that significantly more Luc2^+^/MSCs presented in the aortas of ApoE^-/-^ mice receiving *id*LFA-1-(Ac-G5)-MSCs compared to those treated with MSA-(Ac-G5)-MSCs. Many of Luc^+^/MSCs were enriched in the plaques where ICAM-1 levels are higher (Figure 5A). This is consistent with the levels of ICAM-1 that are robustly expressed in endothelium covering or within plaques. We also observed homing of some Luc2^+^/MSCs to the lumen where there is absence of plaques but ICAM-1 levels are also elevated (Figure 5B). Luc2^+^ *id*LFA-1-(Ac-G5)-MSCs were detectable in the lumen of aorta and small vessels and capillaries in tunica intima and adventitia. In contrast, significantly lower numbers of Luc2^+^ MSA-(Ac-G5)-MSCs were detectable in the aorta of control ApoE^-/-^ mice. No Luc2^+^ cells could be detectable in the aorta of ApoE^-/-^ control mice injected with PBS. Imaging with higher magnification revealed attachment of Luc2^+^/MSCs delivered by *id*LFA-1-(Ac-G5)-nanocarriers to the lumen of aorta. Quantitative data is shown in Figure 5C. Our data demonstrated that *id*LFA-1-(Ac-G5)-nanocarriers can successfully direct circulating MSCs home to inflamed endothelium at atherosclerotic aorta lesions of ApoE^-/-^ mice.

**Figure 5.**
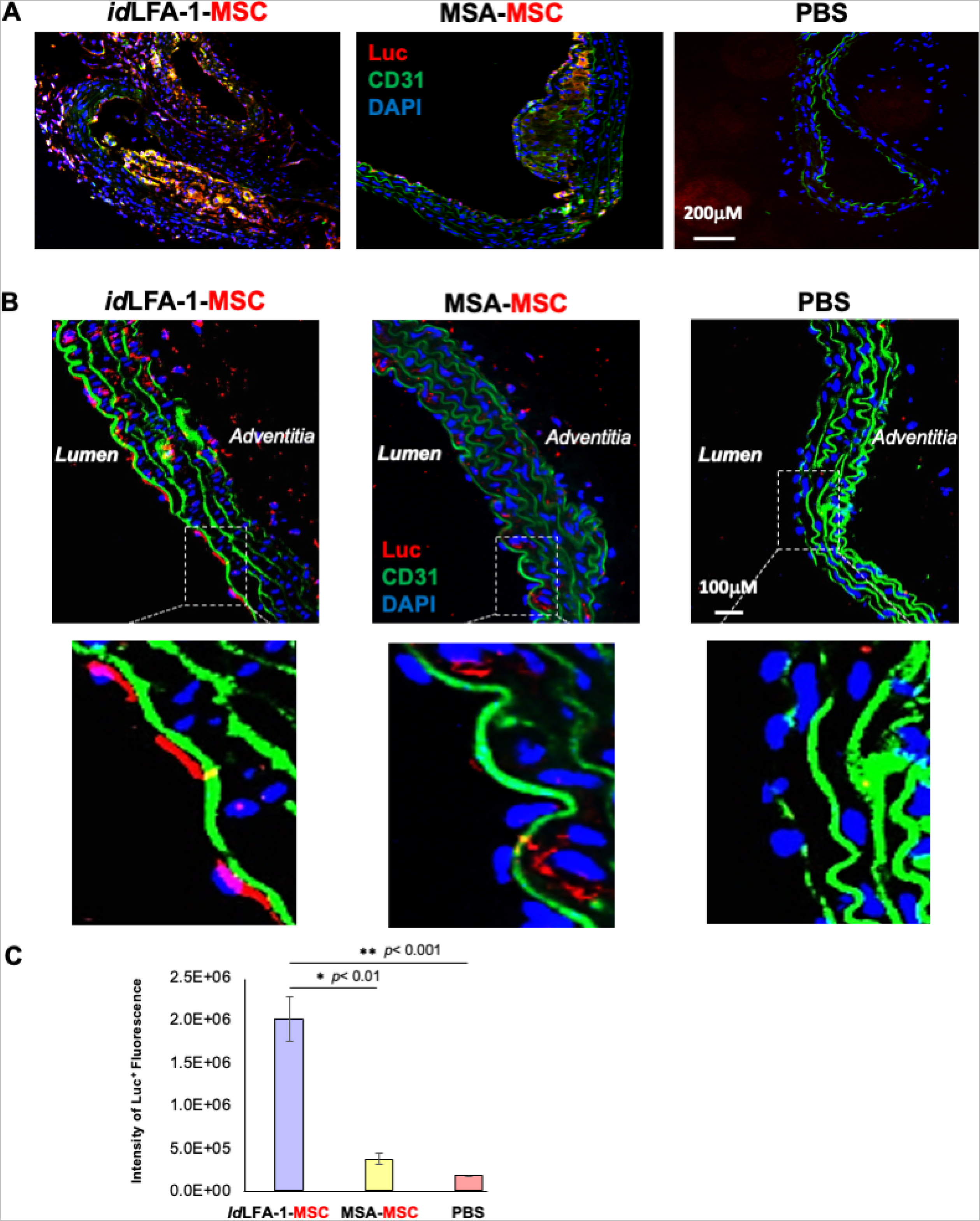
Targeted delivery of *id*LFA-1-nanocarrier-coated MSCs to inflamed aorta in ApoE^-/-^ mice. **A**. Representative immunofluorescent microscopy imaging shows increased colocalization (orange and purple) of *id*LFA-1-coated Luc^+^ MSCs (Red) with nuclei (blue) at aortic plaque sites of aortas harvested from ApoE^-/-^ mice fed a HFD 30 minutes after luciferin injection compared to PBS and murine serum albumin-coated MSC controls. **B**. Representative immunofluorescent microscopy imaging shows increased binding of *id*LFA-1-coated Luc^+^ MSCs (Red) with nuclei (blue) at lumen in non-plaque areas of aortas harvested from ApoE^-/-^ mice fed a HFD 30 minutes after luciferin injection compared to PBS and murine serum albumin-coated MSC controls. Higher magnification images demonstrating Luc^+^ MSCs (Red) attachment at the luminal endothelium (green, along with strong green autofluorescence from elastin). **C**. Quantitative data (ANOVA analysis) of fluorescent images of Luc^+^ MSC signal. Asterisks denote statistical significance (* *p*<0.01, ** *p*<0.001).

### Biodistribution of idLFA-1 and idLFA-1-nanocarrier-coated MSCs in ApoE^-/-^ mouse model

Next, we addressed how specifically *id*LFA-1-Vivo-tag XL680 can traffic and bind on atherosclerotic aorta lesions. Whole-body IVIS scan imaging did not detect that infused *id*LFA-1-Vivo-tag XL680 are enriched in any major organs except in the region of the aortas (see Fig 4B), suggesting that *id*LFA-1 can specifically bind to inflamed endothelium rather than normal endothelium. Immediately following whole-body scanning, the lungs, livers and bladders of the animals were excised and subjected to IVIS scan. No obvious signals were detectable in lungs harvested from 4 groups of ApoE^-/-^ mice injected with *id*LFA-1-Vivo-tag XL680, MSA-Vivo-tag XL680, PBS and Vivo-tag XL680 along, respectively (Figure 6A). Although some Vivo-tag XL680 signal was detected in livers of ApoE^-/-^ mice injected with *id*LFA-1-Vivo-tag XL680 and MSA-Vivo-tag XL680, however, there was no significant difference in the intensity of Vivo-tag XL680 signals between these two groups of livers (Figure 6A, *right*), thus indicating that these signals were not due to specific binding of *id*LFA-1 or MSA to the liver. This may be due in part to partially inflamed endothelium in liver of ApoE^-/-^ mice induced by the HFD.

**Figure 6.**
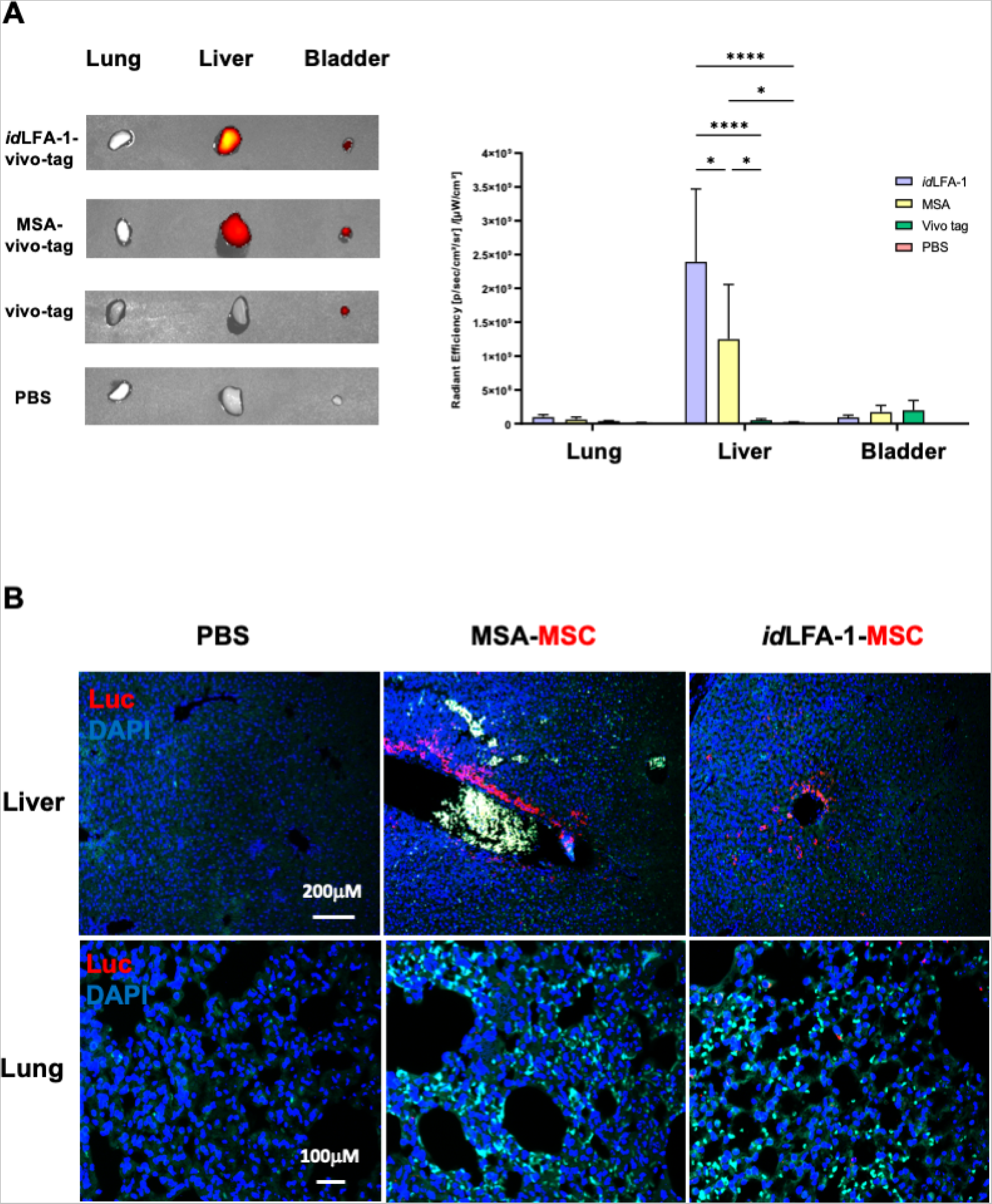
Biodistribution of *id*LFA-1 and *id*LFA-1-nanocarrier-coated MSCs. **A.** *Left*: Fluorescent dye signal intensity showing *id*LFA-1-vivo-tag homing to liver without homing to lung. The vivo-tag signals detectable in the bladder are shown as systemic loading control. *Right*: Quantitative data (ANOVA analysis) of fluorescent images of vivo-tag signals. Asterisks denote statistical significance (* *p*<0.05, **** *p*<0.001), n=5/group. **B**. Representative immunofluorescent microscopy imaging shows comparable deposition of *id*LFA-1-coated Luc^+^ MSCs (Red) with nuclei (blue) at hepatic sinusoids harvested 30 minutes after systemic injection from ApoE^-/-^ mice fed a HFD as that injected with murine serum albumin-coated MSCs. Imaging of tissues from mice injected with PBS are used as baseline controls.

To further test biodistribution of *id*LFA-1-nanocarrier-coated MSCs *vs* MSA-nanocarrier coated MSCs, we conducted immunostaining studies using anti-Luc antibody to examine potential localization of Luc2^+^/MSC coated with of *id*LFA-1-nanocarrier or MSA-nanocarrier in the lung and liver of ApoE^-/-^ mice. Similar as shown in Figure 6A, no obvious Luc2^+^ MSCs were detectable in the lungs (Figure 6B, *bottom*). There were some Luc2^+^ MSCs present in the surroundings of hepatic vessels, however, there was no significant difference between these two groups of livers (Figure 6B, *top*), suggesting that these MSCs are not specifically directed to liver by either the *id*LFA-1-nanocarrier or the MSA-nanocarrier. This may be partially due to the high vessel density in hepatic tissue where infused circulating *id*LFA-1 or MSA can traffic and temporarily locate and enrich or it could be attributed to HFD-induced hepatic inflammation, which could lead to binding of the nanocarrier ot the liver. This data suggests that *id*LFA-1-nanocarriers can selectively direct MSCs to home to the inflamed endothelium at atherosclerotic aorta lesions of ApoE^-/-^ mice.

## 4. DISCUSSION

This work demonstrates an innovative utilization of an adhesion molecule moiety to design nanocarriers for cell surface coating such as to direct the injected cells to home specifically to diseased tissue. This cell delivery system mediated by modified nanocarriers offers superior specificity compared to conventional stem cell delivery with a significantly greater number of cells delivered to target tissues. By exploiting the relationship between a highly or selectively expressed adhesion molecule on luminal endothelial cells and its cognate adhesion moiety in pathological states such as atherosclerosis, therapeutic cells can discriminately traffic to target disease areas upon systemic injection. Here, we identified ICAM-1 to be highly expressed on endothelium in sites with atherosclerotic plaque burden and created nanocarriers with *id*LFA-1 to potentiate MSC delivery to diseased areas. This is the first demonstration of this nanocarrier-based method of targeted delivery to atherosclerotic plaque by employing the highly selective interaction between *id*LFA-1 ICAM-1. It should be noted that interactions between ligand and receptor can result in unintended effects including signal transduction and pro-inflammatory stimulation in endothelial cells. Thus, it is critical to carefully select specific moieties/adhesion domains for complexation with nanocarriers to result in maximum binding capacity without complete induction of unwanted intracellular cell signaling. In that regard, our method to install specific moieties/adhesion domains on the cell surface via nanocarrier can avoid potential unintended consequence of intracellular cell signaling initiated by interactions between ligand and receptor, which may induce uncontrolled cell differentiation, particularly when infusing MSCs and any other types of stem/progenitor cells.

Our nanocarrier-coated cell delivery technology platform employs the anionic properties of the cell membrane to associate with nanocarriers composed of cationic dendrimers. This ionic bond results in cell-surface electrostatic attachment of nanocarrier-dendrimer complexes via complexation. Given the reversible, non-covalent relationship of this complexation, nanocarriers may theoretically disassociate with MSCs after injection and bond with other cell types *in vivo*. Despite this, our data indicates that nanocarrier-coated cells administered systemically maintain the ability to selectively migrate to target tissues thereby demonstrating the residual high fidelity of these nanocarrier complexes on MSCs after injection. Other associations between nanocarriers and cell surface substrates such as protein or carbohydrates as well as incomplete internalization of nanocarriers after initial coalescence of nanocarriers on the cell membrane could also explain this observed phenomenon. Another benefit of dendrimer-nanocarrier cell membrane coating is that eventual complete internalization of these complexes by cells allows for non-toxic elimination from the body and, subsequently, result in enhanced biocompatibility. Given that this platform relies on physical non-covalent interactions of adhesion molecules on the cell surface without manipulation of cell populations underlying genetic expression or protein machinery, it can be tailored to incorporate nearly any adhesion moiety/ligand on any cell to program targeted delivery. For example, we have previously demonstrated the role of an inducible cell adhesion molecule, E-selectin, in mediating endothelial progenitor cell recruitment to wound sites.^5^ By cell surface coating of MSCs with Ac-G5-sE-sel dendrimer nanocarriers, systemic cell delivery of MSCs to wounded tissue was substantially increased. This resulted in direct biological responses through both accelerated rates of wound closure and pro-angiogenic effects of MSCs through enhanced neovascularization in wound bed tissue.^5^

Importantly, we further demonstrated the efficacy of these nanocarriers in enhancing *id*LFA-1/ICAM-1 interactions at three different levels: (1) *id*LFA-1/ICAM-1 *in vitro* protein-EC cell binding, (2) *id*LFA-1-Cy5-(Ac-G5)/ICAM-1 nanocarrier-EC cell binding *in vitro*, and (3) MSC-EC binding via *id*LFA-1-Cy5-(Ac-G5)/ICAM-1 both *in vitro* and *in vivo*. Previous studies demonstrated the utility of *id*LFA-1 as a strategy to deliver drug and/or target molecules to different cell types expressing surface ICAM-1.^24,34^ Others further demonstrated the ability of modified nanoparticles in delivering drug cargo to ICAM-1 expressing cell types for localization to target cell phenotypes.^34^ However, manipulation of nanocarrier particles with *id*LFA-1 to improve delivery of cell therapy has not been previously investigated. Given the specific homing of coated-MSCs to luminal endothelium in atherosclerotic lesions, this suggests that ICAM-1 is present at high enough levels to recruit circulating cells at disease sites, even in locations subjected to the high velocity/dynamic circulatory flow of the aorta. Furthermore, the fact that BSA-nanocarrier coated-MSCs achieved only partial localization to sites of atherosclerosis demonstrates that upregulation of ICAM-1 levels alone are not sufficient to recruit unmodified MSCs into the aorta. Given that nanocarrier-coated MSCs were administered systemically, it is conceivable that these cells could also traffic via capillaries and postcapillary venules to alternative disease sites at peripheral locations that also express the target binding molecule of interest. As with any systemic administration method, there is the potential for indiscriminate trafficking to other organs. In particular, the lung and liver can decrease the effectiveness of cell engraftment given their dense capillary networks, which typically persist as cell filtration obstacles to conventional MSC therapies infused via venous routes.^1^ We specifically examined the lung and liver, two major organs which act as filter sinks collecting blood from systemic circulation veins (lung) and arteries (liver), and bladders that collect/store expelling *id*LFA-1 (*id*LFA-1-Vivo-Tag XL680), thus serving as loading control. Interestingly, there was no significant signal intensity detected in *ex vivo* lung tissue. However, there was partial colocalization of *id*LFA-1-coated MSCs to hepatic vessels after systemic administration in our study. This can be caused in part by hepatic lipid elevation-induced inflammation, which is well-known to occur in ApoE^-/-^ mice fed with HFD.^35,36^ Excessive intake of fat can result in hepatic lipid accumulation, thereby leading to non-alcoholic fatty liver disease (NAFLD). NAFLD livers exhibit inflammation due to infiltration of inflammatory macrophages and other myeloid cells in the hepatic parenchymal area.^37^ NAFLD-induced hepatic inflammation can further result in upregulation of ICAM-1 on the endothelium within fatty liver tissues.^38^ Despite partial deposition of MSCs in liver, there was still a significantly greater amount of *id*LFA-1-coated MSCs delivered to inflamed aortic endothelium. In addition to NAFLD-induced hepatic inflammation, partial deposition of MSCs in liver could also be ascribed to hepatic clearance, which is known to occur in other MSC therapies.^1,39^ Compared to biodistribution in hepatic tissue, we observed similar amounts of MSA-nanocarrier-coated MSCs present in the surrounding area of hepatic vessels/sinusoids. This may suggest that deposition of MSCs in liver can be due to a general indiscriminate clearance of MSC therapy from the body.

MSC therapy has shown promise in preclinical studies for reducing atherosclerotic burden and plaque progression given their numerous beneficial immunomodulatory effects. Transplantation of MSCs is associated with an increase in anti-inflammatory cytokines such as IL-10 and TGF-β, as well as attenuation of the release of inflammatory mediators such as TNF-α and NF-κB.^40–43^ A limitation of this work was the inability to examine the therapeutic effect of MSC therapy upon atherosclerotic lesions as our construct employed the human *id*LFA-1, which is immunogenic to experimental mice. Subsequent work should take into consideration the design and homology of target proteins of interest in inducing maximum immunotolerance across preclinical *in vivo* models and translational clinical studies. An advantage of the described method is the ability to customize nanocarriers to enable direct delivery of various cell types to different tissue types, thus highlighting the significant potential as a versatile platform for cell-based therapeutics for a myriad of diseases. Finally, this approach can have broad utility and applications for cell-based therapy in transplantation, cardiovascular disease, autoimmune pathologies, and regenerative medicine.

## 5. CONCLUSIONS

We describe a novel application of newly-designed nanocarriers for cell surface coating to guide the homing of systemic administrated circulating cells to disease sites. Specific cell adhesion moieties/molecule tailored nanocarriers are able to act as a “GPS” and guide coated cells to find destination via targeting the blood/vessel interface for cell delivery. The ability to customize this nanocarrier technology based on specific adhesion moieties/molecule and their cognate binding counterpart highly or selectively expressed on the endothelium in diseased tissue enables this as a highly versatile platform to enhance systemic delivery of cell-based therapeutics for a broad variety of clinically relevant translational purposes.

## ACKNOWLEDGMENTS

We thank the University of Miami Analytical Imaging Core Facility for assistance with IVIS Imaging and data analysis.

## Author Contributions

Study Conception and Design: ED, SKD, SD, ZJL, OCV. Acquisition of Data: All authors except TJS. Analysis and Interpretation of Data: All authors except TJS. Drafting of Manuscript: CTH, ED, SKD, SD, ZJL, OCV.

